# mtFociCounter for automated single-cell mitochondrial nucleoid quantification and reproducible foci analysis

**DOI:** 10.1101/2022.08.13.503663

**Authors:** Timo Rey, Luis Carlos Tábara, Julien Prudent, Michal Minczuk

## Abstract

Mitochondrial DNA (mtDNA) encodes the core subunits for OXPHOS and is essential in eukaryotes. mtDNA is packed into distinct foci (nucleoids) inside mitochondria, and the number of mtDNA differs between cell-types, and is affected in several human diseases. Today, common protocols estimate per-cell mtDNA-molecule numbers by sequencing or qPCR from bulk samples. However, this does not allow insight into cell-to-cell heterogeneity and can mask phenotypical sub-populations. Here, we present *mtFociCounter*, a single-cell image-analysis tool for reproducible quantification of nucleoids and other foci. *mtFociCounter* is a light-weight, open-source freeware and overcomes current limitations to reproducible single-cell analysis of mitochondrial foci. We demonstrate its use by analysing 2165 single fibroblasts, and observe a large cell-to-cell heterogeneity in nucleoid numbers. In addition, *mtFociCounter* quantifies mitochondrial content and our results show good correlation (R=0.90) between nucleoid number and mitochondrial area, and we find nucleoid density is less variable than nucleoid numbers in wild-type cells. Finally, we demonstrate *mtFociCounter* readily detects differences in foci numbers upon sample-treatment, and applies to superresolution microscopy. Together, we present *mtFociCounter* as a solution to reproducibly quantify cellular foci in single cells and our results highlight the importance of accounting for cell-to-cell variance and mitochondrial context in mitochondrial nucleoid analysis.

## Introduction

Mitochondria are dynamic and heterogeneous cellular organelles. They occur in a vast range of forms from small bean-shapes to large tubular networks, depending on the state and type of their host cell [1]. To build the oxidative phosphorylation (OXPHOS) apparatus, crucial for ATP production, mitochondria maintain express their own genome, mitochondrial (mt)DNA. Mammalian mtDNA is circular, encodes 37 essential genes on ∼16.6kb and is stored in small (∼100nm) nucleoprotein-granules, termed nucleoids [2,3]. The amount of mtDNA is tightly regulated at the cellular level and ranges from 100 to 100’000 copies, depending on cell state and type [4–7]. Aberrations of mtDNA copy-number control are associated with mitochondrial disorders and cardiovascular disease, cancer and neurodegeneration [6,8–10]. Mitochondrial Transcription Factor A (TFAM) has been identified as a key factor to regulate mtDNA transcription, replication and compaction [11–13] and downregulation of TFAM decreases mtDNA copy number in cultured cells [12,14]. However, the mechanisms by which healthy mammalian cells regulate mtDNA and nucleoid numbers are still not understood [9]. Reliable quantification of the number of nucleoids, therefore, remains crucial to elucidate the mechanisms of cellular energy homeostasis in health and disease.

To determine the number of mtDNA molecules per cell, current methods rely on quantitative PCR and sequencing [6]. Hereby, mtDNA from bulk samples is normalised to nuclear DNA to obtain per-cell estimates. Yet, these methods do not preserve information beyond the average number of mtDNA molecules from millions of cells and do not reveal cell-to-cell heterogeneity. The recent droplet digital PCR approach allows measurement of mtDNA copy-numbers in single-cells but is limited in throughput and and looses all contextual information, as it requires mtDNA extraction [15,16].

Mitochondrial nucleoids and other foci, such as mitochondrial RNA granules (MRG), can also be visualised by fluorescence microscopy [17–20]. The number of nucleoids per cell can therefore be counted directly from live or fixed cell images, allowing to preserve context [7,21]. In addition, quantitative methods for image analysis developed for one foci-type are readily transferable to other types, allowing direct comparison, for instance between one nucleoid marker and another [2]. Currently, mitochondrial nucleoids are typically counted manually or manual thresholding, impacting reproducibility and constraining the throughput to few 10s of cells and 100s of foci [22,23]. Automation of foci-counting drastically reduces random errors and observer biases and allows scaling to 100s of cells and 10’000s of foci [3,24,25]. However, previous reports for automated mitochondrial nucleoid counting do not disclose the analysis code and the choice of influential parameters, restricting reproducibility and comparison of results.

Here, we present an easy-to-use, free and open source software tool to quantify mitochondrial foci in single cells, which we named Mitochondrial Foci Counter or *mtFociCounter. mtFociCounter* provides a 100% reproducible analysis, from raw image to final results, and allows transparent parameter choice. We showcase the use of *mtFociCounter* by analysing more than 2’000 single cells and over 500’000 foci from confocal and superresolution fluorescence microscopy, and find a very large heterogeneity of mitochondrial nucleoid numbers across single cells. In addition to foci-counts, *mtFociCounter* allows to estimate the total area of the mitochondrial network in each cell. This feature allows to investigate the relationship between nucleoid numbers and mitochondrial area, for which we find a good correlation. In further assessment of *mtFociCounter* capabilities, we not only validate its transferability to an additional, human cell model but also detect and quantify decreased nucleoid numbers upon TFAM knockdown, in U2OS cells.

Taken together, *mtFociCounter* provides a novel tool for reproducible single-cell analysis of mitochondrial foci, and its modular design and open source code enable easy adaptation to specific needs, retaining a uniform output format between studies. *mtFociCounter* allows to quantify nucleoid numbers in single cells at high throughput and to study mtDNA abundance in health and disease.

### Material and methods

#### Cell culture and transfections

NIH/3t3 fibroblast cells (3t3) were obtained from American Type Culture Collection (ATCC) (ATCC: CRL-1658™) and cultured in growth medium with DMEM GlutaMax (Gibco: 31966-021) supplemented with 1% PenStrep and 10% Calf Serum (ATCC: 30-2030). Calf Serum instead of Fetal Calf Serum is used to maintain proliferative cell populations of 3t3 fibroblasts. Stable cell-lines expressing MTS-dsRed were created as previously described [26]. In brief, cells were transformed with lentivirus to express MTS-dsRed, whereby sufficiently high expression-levels were achieved. To ensure genetic coherence across experiments, cells were cultured for no more than 20 passages before thawing a fresh cryo-vial. U2OS cells were obtained from ATCC and cultured in McCoy’s medium (Life Technologies, Gibco®) supplemented with 10% fetal bovine serum (Life Technologies, Gibco®), 2 mM L-glutamine, non-essential amino acids and 100 units/mL penicillin/streptomycin (all GIBCO). All cell-cultures were incubated at t 37°C and 5% CO_2_.

For small interference RNA experiments, U2OS were transfected using Lipofectamine RNAi max (Invitrogen) with 20 nM siRNA for three days following the manufacturer’s instructions. siRNA used, were: Non-targeting siRNA (ON-TARGETplus SMARTpool D-0001810-10-20) and siTFAM (ON-TARGETplus Human TFAM (7019) siRNA - SMARTpool, L-019734-00-0005).

### Immunofluorescence

3t3 cells were seeded on 12 mm glass slide cover slips (Epredia: CB00130RAC20MNZ0) in different wells of 24-well plates, and at a density of 50’000 cells per well. The next day, the medium was aspirated and cells were fixed with 300 μL of pre-warmed 4% PFA (Thermo scientific: J19943-K2) for 20 min at room temperature (RT). Samples were then rinsed with PBS once, and washed with PBS two times for 10 min at RT. Occasionally, samples were refrigerated (4 °C) for few hours at this step, before the process was continued later on the same day. The cells were then permeabilised for 10-20 min with 250uL of 0.1% Triton X-100 (Fluka: 93420), before blocking with 300 μL of 5% BSA (Sigma: A7030-100G) in PBS for 15-30 min at RT. For immunostaining, anti-DNA antibodies (EMD Millipore: CBL186) and anti-TOMM20 antibodies (Proteintech: 11802-1-AP) were then diluted in 5% BSA in PBS at a ratio of 1:250 (a solution of at least 125 μL was made to reduce pipetting errors), and cover slips were flipped onto 20 μL drops of antibody-solution on parafilm for 1 hour of incubation in a humid chamber at RT. Samples were then rinsed with PBS once and washed two times for 5-15 min with PBS. Next, anti-IgM Alexa Fluor 647 antibodies (Invitrogen: A21238) were diluted in 5% BSA at a ratio of 1:750 (a solution of at least 350 μL was made) and cover slips incubated as described above, for 45 min. Where applicable, Wheat Germ Agglutinin Alexa Fluor 488 (ThermoFischer: W11261) was also added. Cover slips were then flipped onto drops of Hoechst 33342 (Invitrogen: 10337) diluted in PBS (1:5000 or 1:2000) and incubated for 15 min before again rinsing and washing in PBS. Individual cover slips were then mounted onto a drop of ProLong Gold (Invitrogen: P10144) and cured at 4 °C over night, in a dark but slightly opened chamber. Samples were always imaged the next day, to avoid sample deterioration.

U2OS cells were fixed in 5 % PFA in PBS at 37°C for 15 min, then washed three times with PBS, followed by incubation with 50 mM ammonium chloride in PBS to quench the unspecific fluorescence signal from aldehyde groups and three additional washes in PBS were performed. Cells were then permeabilized in 0.1 % Triton X-100 in PBS for 10 min before blocking in 10% FBS in PBS for 20 min, followed by incubation with anti-TOMM20 (Proteintech: 11802-1-AP) and anti-DNA (EMD Millipore: CBL186) primary antibodies in 5% FBS in PBS, for 2 h at RT. After 3 washes with 5 % FBS in PBS, cells were incubated with goat anti-Rabbit IgG (H+L) Alexa Fluor 488 conjugate (A-11070) and goat anti-Mouse IgM (Heavy chain) cross-adsorbed Alexa Fluor 568 conjugate (A-21043) secondary antibodies for 1 h at RT. After 3 washes in PBS, coverslips were mounted onto slides using Dako fluorescence mounting medium (Dako).

### Microscopy

All spinning disk confocal images were acquired using an Andor Dragonfly 500 system on a Nikon Eclipse TiE inverted microscope. Image stacks were acquired with a 100x or 60x objective lens (NA1.4). We used 405 nm, 568 nm or 633 nm excitation lasers and acquired images on a Zyla 4.2 PLUS sCMOS camera with Fusion software (Andor). Laser intensity and exposure times were optimised for wild-type or control cells of each cell type, and kept constant across all acquisitions. Images for z-stacks of 3t3 cells were acquired bottom to top every 0.3 μm spanning the entire cell or every 0.2 μm for U2OS comprising 7 images. Superresolution microscopy images were acquired on a Zeiss Elyra7 microscope with a Lattice SIM^2^ processing system. This comprises a Zeiss AxioObserver 7 Inverted Microscope with a Zeiss Elyra7 imaging head, environmental chamber, dual cameras with DuoLink, and four lasers including a Solid-State Diode laser with 50mW maximum power emission at wavelength 405nm, two OPSL continuous wave lasers with maximum power of 500mW at emission wavelength 488nm and 561nm respectively, and a Solid-State Diode continuous wave of 500W at 643nm emission. All z-stack images were acquired with 0.126μm step-size and 1280 by 1280 pixels (0.063μm by 0.063μm) using a Plan-Apochromat 63x/1.4 Oil DIC M27 objective. The three channels were acquired sequentially with 13 phase images at 50ms exposure and 50% (405nm), 5% (642nm) or 5-15% (561nm) laser power, whereby Hoechst was acquired with Band Pass filters BP 420-480 and BP 495-550, whereas dsRed and AlexaFluor 647 were acquired with BP 570-620 and LP 655 filters, on Dual PCO.Edge 4.2 sCMOS cameras with a DuoLink adaptor.

### Image Analysis

All image processing and analysis was done using the *mtFociCounter* workflow described in this article. We set the prominence range for our images between 5 and 200 au. For 3t3 samples imaged by confocal, where the Hoechst-signal was too low, as well as for superresolution images, where nuclei are too well resolved, we used manual segmentation for nuclei using the Hoechst-channel as input. For U2OS samples, we used the mitochondria channel to manually segment the nuclear area. Mitochondria were automatically detected, and used to filter the foci, as well as to determine the mitochondrial area. The raw-data was then processed with python 3 and jupyter notebooks. *mtFociCounter* and all jupyter notebooks are available on Github (Github.com/TimoHenry).

### Statistical Analysis

Data-Normality was tested by distribution assessment, Q-Q plots and by CDF-distribution analysis and Kolmogorov-smirnov testing against simulated data-sets (see Supplementary Data). Subsequently, non-parametric tests were used where non-normal datasets were compared. ASpecific alpha-values and statistical tests which were used are indicated for each figure.ll statistical analysis was performed in python 3 and exact algorithms and package versions are provided in the corresponding jupyter notebooks (Github.com/TimoHenry).

### Software

Only free and open-source software was used for this project, with the exception of microscope-steering, as indicated above. For analysis, MiniConda was used as a package manager to run python 3 and jupyter notebooks. For writing, reference management and figure-generation, LibreOffice, Zotero, Fiji (ImageJ) [27] and Inkscape were used respectively.

### Results

#### mtFociCounter allows single cell analysis of mitochondrial foci

The overarching workflow to quantify mitochondrial foci in single cells by *mtFociCounter* is represented in **Figure 1**. *mtFociCounter* is a Fiji-plugin composed of three modules: *ToTiff, CellOutline* and *FociCounter* (**Supplementary Figure 1**). Each module can be run independently, and is discussed briefly in the following.

**Figure 1.**
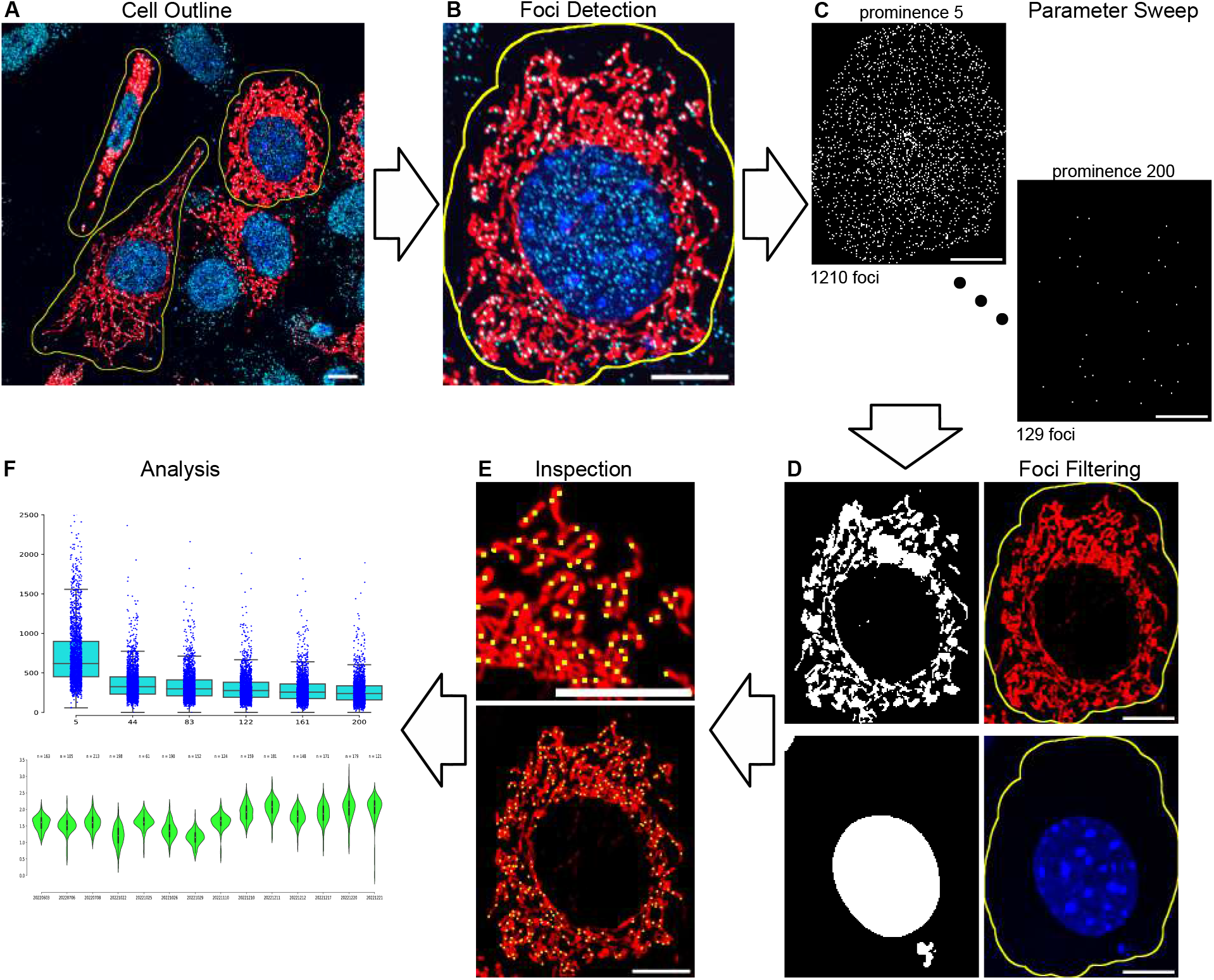
Workflow of Mitochondrial Foci Counter (*mtFociCounter*). (**A**) Exemplary Spinning Disk Confocal image of 3t3 fibroblasts which express MTS-dsRed (red), used as input for *mtFociCounter*. Mitochondrial nucleoids are stained with primary antibodies against dsDNA and AF647 secondary antibodies (cyan), and nuclei are stained by Hoechst 33342 (blue). Manually drawn segmentation lines are shown in yellow. (**B**) One individual segmented cell from **A**, as output by *CellOutline* module. (**C**) Two of six binary images produced by *FociCounter* module performing a parameter sweep between prominence values 5 and 200 au on the image in **B**. The number of detected mitochondrial foci is indicated below each image. (**D**) Automatically generated binary masks which are used to filter detected foci from **C**. (**E**) Bottom shows detected foci (prominence = 83) after filtration (yellow), overlaid onto mitochondria (red) from image **B**. Top shows zoomed-in image from bottom. (**F**) Representative analytical graphs for the parameter sweep (top) or comparison of 14 independent experimental replicates (bottom). All scale bars represent 10μm. Image intensities were adjusted manually for representation, but not for quantitative analysis.

The first module, *ToTiff* converts images with microscope-specific formats into TIFF-files. The user can choose a directory with input files and TIFF files are saved in a new folder. We then consider the TIFF-files as raw data for analysis, storage and upload to public databases, and delete original (non-TIFF) image files upon conversion, to avoid data-multiplication.

The second module, *CellOutline* uses TIFF-files as input and allows segmentation of individual cells (**Figure 1A**). For multi-colour images, combined channels are displayed and 3 dimensional images are z-projected using maximum intensity by default. We then select individual cells by manual segmentation of a region of interest (ROI). All ROIs are saved to ensure perfect reproducibility of the selection before a new image is created for every selected cell (**Figure 1B**). Manual segmentation allows to exclude debris or overlapping cells, and ensures high-quality input data (**Supplementary Figure 2**). Depending on the experimental set-up, the modular design of *mtFociCounter* allows cell-segmentation by other workflows, such as CellCatcher [28] or CellProfiler [29], to be used as input to the last module.

**Figure 2.**
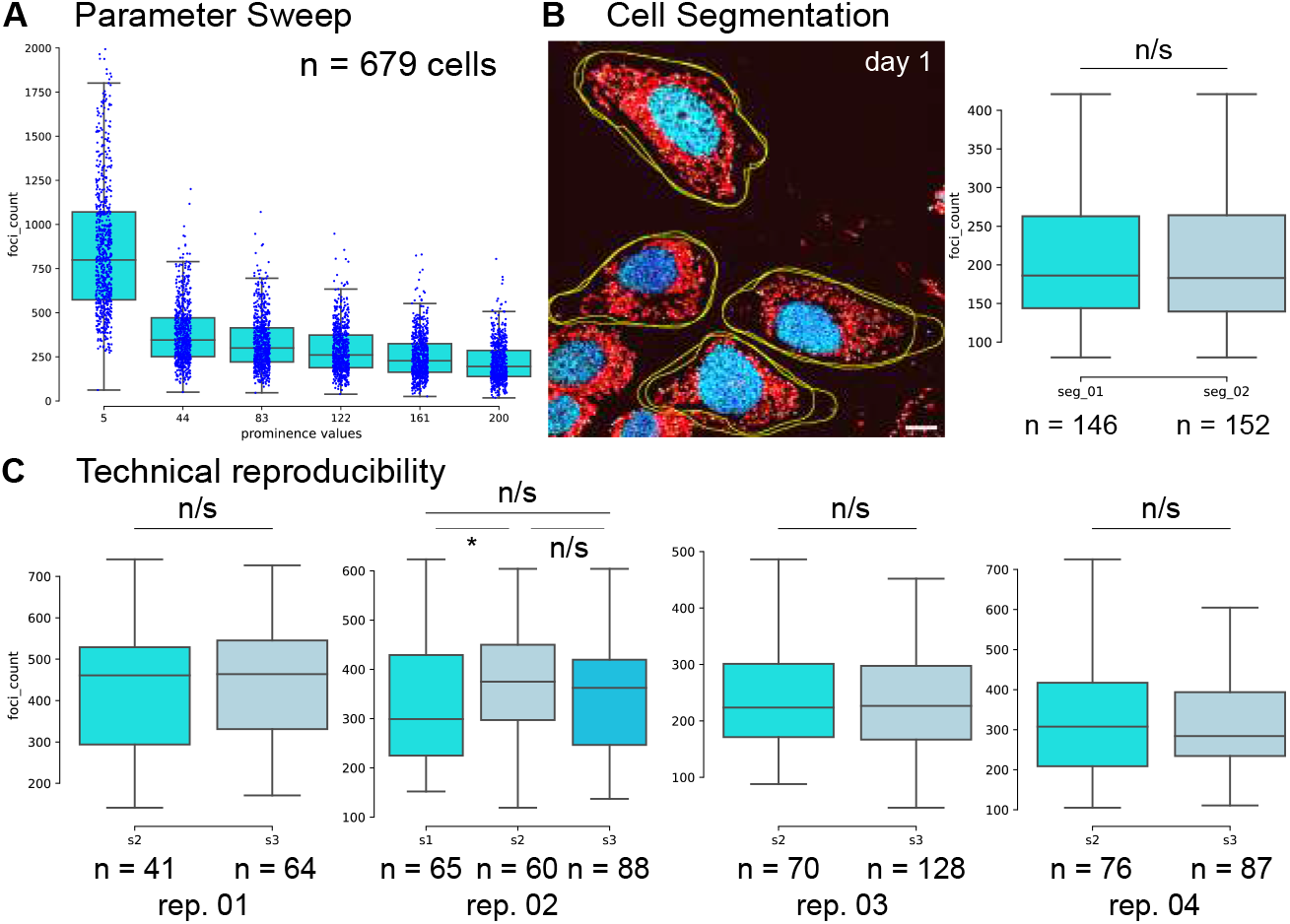
Experimental verification and parameter optimisation by *mtFociCounter*. (**A**) Histogram plots of the number of detected foci (nucleoids) in 3t3 WT cells (n=679 single cells) as a function of 6 different prominence values demonstrates the use of a parameter-sweep. Blue dots represent single-cell measurements. (**B**) Representative FOV (left) with the outlines (yellow) from two separate manual segmentations of four individual cells, and comparison of the number of nucleoids detected (right) from the same sample after separate rounds of cell segmentation (n=146 analysed cells after first segmentation and n=152 analysed cells after second segmentation). (**C**) Comparison of the detected number of nucleoids from two or three technical replicates, with samples processed in parallel in four independent experiments (processed on different days). All box plots represent the median and the first and third quartile. Whiskers represent the rest of the distribution except outliers. Two-sided Kolmogorov-Smirnoff tests were used with alpha = 5% and * denoting p<0.05. Scale bar is 10μm.

The third module, *FociCounter* quantifies the number of foci in segmented single cells. First, 2D TIFF-images with an ROI are loaded (**Figure 1B**). Individual foci are then detected by the *FindMaxima* algorithm and a single parameter, termed ‘prominence’. *FociCounter* encourages to perform a parameter sweep of six prominence values and requires to define a range of values (**Figure 1C**). Detected maxima are counted and binary images of detected foci as well as a list of ROIs are saved for future inspection. Optionally, *FociCounter* allows to filter the detected foci (**Figure 1D**). To ensure only mitochondrial foci are counted, we enable the “Filter with mitochondria” option and use automated segmentation to create a mitochondrial mask. Again, a gateway for pre-segmented masks allows the use of external software such as Ilastik [30] or Weka [31] for segmentation, and manual segmentation is possible. The area of the inclusion mask (mitochondrial mask) is measured for every cell and saved as an output [32]. Because nuclear DNA can produce false positive mtDNA-signals in mitochondria which overlap with the nucleus after z-projection, an option “Subtract nucleus” allows to filter mitochondrial foci by exclusion of the nuclear volume (**Figure 1D**). We use Hoechst 33342 to stain the nucleus for automated segmentation by *FociCounter*. Alternatively, images of mitochondria can be used for manual segmentation, as the nuclear shape is identifiable by eye. Binary images of segmented nuclei and mitochondria are saved for every cell to ensure perfect traceability and reproducibility of the complete analysis pipeline. After filtering, the quality foci detection can be visually verified by overlay of every detected foci onto the corresponding image (**Figure 1E**). Finally, a single CSV-file is generated with number of foci and total mitochondrial network area per cell, for further analysis (**Figure 1F**).

### Nucleoid analysis by mtFociCounter in 3t3 fibroblasts

To demonstrate the use of *mtFociCounter* for mitochondrial foci quantification we analysed the number of mitochondrial nucleoids in wild-type (WT) 3t3 mouse fibroblast cells. We found clean antibody-staining of mitochondria in 3t3 mouse cells challenging (**Supplementary Figure 2A**), and therefore created a stable cell line expressing mitochondrial matrix targeted fluorophores (MTS-dsRed), as previously described [26]. Cells were then fixed and nucleoids stained with antibodies against dsDNA, which highlights mitochondrial nucleoids. The nucleus was stained with Hoechst 33342 and three-colour fluorescence images were acquired on a standard spinning disk confocal microscope (**Figure 1A**). To convert raw images from *.ims to *.tiff files the *ToTiff* was used module. Next, *CellOutline* with default options for maximum-intensity projection and manual segmentation was used to extract individual cells. We observed that 3t3 fibroblasts often grow on top of each other, even at medium and low cell density (**Supplementary Figure 2B**), and that MTS-dsRed is expressed at different levels in individual cells. Overlapping cells with ambiguous boundaries, cells with faint mitochondrial signal as well as dividing cells with condensed chromosomes were, not selected for further quantification (**Supplementary Figure 2C**). To measure the number of mitochondrial nucleoids we next applied *FociCounter* module, filtering out non detection of nuclei and mitochondria, as described above.

### Parameter-sweep promotes analysis of parameter-choice

To test the effect of different prominence values on the detected number of foci *mtFociCounter* performs a fully automated parameter-sweep. In our experiments, a low prominence of 5 arbitrary units (au) leads to the detection of a large number of non-specific foci, whereas with a prominence of 200 au many true positive nucleoids are missed (**Figure 1C**). The optimal prominence depends on imaging acquisition and processing parameters and needs to be defined for every experimental workflow. We therefore recommend to begin with coarse sweeps for new projects. In our experiments, we observe a sharp drop of detected foci between 5 and 44au (**Figure 2A**), indicating a clear difference between real positive and false positive signals, in accordance with visually inspected overlays. To validate that the detected foci are indeed nucleoids, we performed control experiments in 3t3 WT cells without dsDNA primary antibodies (**Supplementary Figure 2D**), and with 3t3 cells devoid of mtDNA (rho0 cells) (**Supplementary Figure 2E**). These experiments confirmed that a parameter sweep is necessary but sufficient to identify reliable settings for true positive foci-detection. In the here described work, we fixed the prominence at 83 au for all analysis of 3t3 cells, unless indicated otherwise.53 nucleoids0 nucleoids 299 nucleoids

### mtFociCounter enables analysis of experimental reproducibility

Next, we used the *CellOutline* feature of *mtFociCounter* to assess the observer-bias by repeating manual cell selection on the same images, for two data-sets. We found negligible differences between repetitions of manual cell segmentation (p_1_= 1.000 and p_2_= 0.974 from Two-sample Kolmogorov-Smirnoff Test) (**Figure 2B** and **Supplementary Figure 2F**) and that manual cell segmentation by a trained experimenter is robust and reproducible. Furthermore, we compared four technical duplicates and triplicates and found no significant difference between samples which were processed and imaged in parallel, on the same day (p_1_=0.285, p_2_=0.735, p_3_=0.349 and p_4_=0.982, Two-sample Kolmogorov-Smirnoff Test) except for one sample, which is on the virtue of significance (p_5_=0.025 and 0.055) (**Figure 2C**). Furthermore, we found that the distribution of the number of mitochondrial nucleoids per cell is significantly different between samples which were prepared on different days (p=1.379*10^−29^, Kruskal-Wallis test) (**Supplementary Figure 2G**). Together, this illustrates that the sampling approach is robust but significant variability between independent experiments exists. This indicates that samples which will be compared need to be prepared in parallel and that a sufficiently large sample size (>60 cells per replicate) is preferable.

### Nucleoid numbers in single 3t3 cells are highly variable

To investigate the biological variance of nucleoid numbers between single cells we measure more than 700’000 nucleoids from 2165 single 3t3 WT cells by *mtFociCounter* (**Figure 3A** and **3D**). We observe a mean number of 339 nucleoids (+/-188 std) and a median number of 299 nucleoids (Interquartile range of 201 nucleoids) per single 3t3 WT cell (n=2165 cells). Interestingly, we find the number of nucleoids in 3t3 fibroblasts is not Normally distributed (p=5.674*10^−10^;Two-sample Kolmogorov-Smirnoff Test)(**Supplementary Figure 3A, D, G** and **Figure 3A**). We also compute the mitochondrial area per cell and find a mean of 211.4μm^2^ (+/-137.1μm^2^ std; n=2165 cells). Again, the distribution of mitochondrial area per cell does not follow a Normal distribution (p=1.118*10^−24^); Two-sample Kolmogorov-Smirnoff Test) (**Supplementary Figure 3D, E, H**) and the median mitochondrial area observed is 176.3μm ^2^ with an inter quartile range of 116.9μm^2^(**Figure 3B** and **E**). Together, we thus observe a very wide range of number of nucleoids as well as mitochondrial area in single 3t3 WT cells, with a coefficient of variation of 55.5% for nucleoids per cell and 64.9% for mitochondrial area. These results are in good agreement with two recent reports of mtDNA copy-numbers from single-cell digital droplet PCR [15,16], and overall expected single-cell heterogeneity [33]. Mixed populations of proliferative cells do not necessarily follow a Normal distribution of cell-states, and our observations indicate that determining a population-averaged mean number of nucleoids per cell may mask subtle differences in the underlying single-cell state distribution.

**Figure 3.**
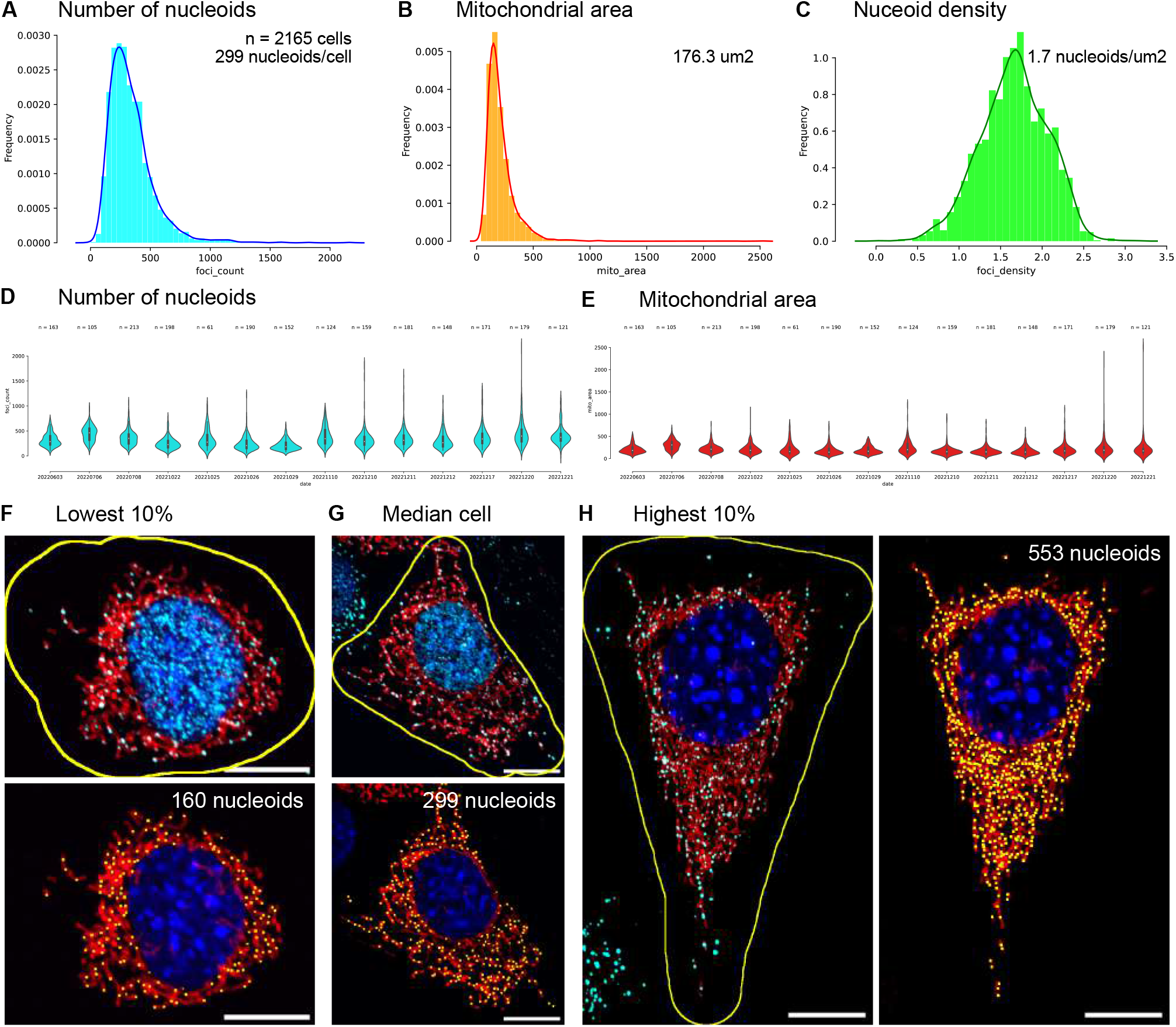
Quantitative analysis of 2165 3t3 WT mouse fibroblasts from Spinning Disk Confocal microscopy by *mtFociCounter*. Histogram and Kernel Density Estimation plots of (**A**) the number of nucleoids, (**B**) the mitochondrial network area in μm^2^ and (**C**) the nucleoid density (number of nucleoids per mitochondrial area) per single cell are shown. A total of 2165 cells from 14 independent experiments were analysed. Median values are indicated. Violin plots for each experiment from **A-C** are shown for the number of nucleoids (**D**) and the mitochondrial are (**E**). Individual number of cells analysed for each sampling date, median and first and third quartiles are indicated. Median single-cell nucleoid numbers are Normally distributed (**Supplementary Figure 3K**). (**F**) Images of the cell denoting the lowest 10% according to the number of detected nucleoids (160), with mitochondria (red), nucleus (blue) and DNA-foci (cyan) and cell-segmentation outline (yellow line) on top or detected foci (yellow boxes) in bottom. (**G**) Images of the cell with the median number of nucleoids (299), and rest as above. (**H**) Images of the cell comprising the 10% highest number of detected nucleoids (553), as above, with anti-DNA (cyan) and cell segmentation (yellow) left, and detected foci right. A prominence level of 82 was used for these analyses. All scale bars are 10μm.

### Nucleoid numbers correlate with mitochondrial network area in single cells

Because the distributions of nucleoid numbers and mitochondrial area resemble each other, we next asked whether these features are related within individual cells. Indeed, we found the number of nucleoids is strongly correlated with the mitochondrial area in 3t3 cells (Corr=0.81, n=2165 cells, **Supplementary Figure 3J**). We then calculated a nucleoid density by normalising the absolute number of nucleoids with the respective mitochondrial area and found a single-cell mean nucleoid density of 1.7 nucleoids/μm^2^ (+/-0.4 std; n=2165 cells) (**Figure C**). Interestingly, this nucleoid density is Normally distributed (**Supplementary Figure 3C, F, I**, p=0.112; Two-sample Kolmogorov-Smirnoff Test) and the coefficient of variation is substantially smaller at 24.1%. Together, cells with fewer nucleoids are thus smaller whereas larger cells tend to have more nucleoids (**Figure 3F-H**). This is in accordance with literature that found a semi-regulated spacing of mitochondrial nucleoids across eukaryotic species [34,35], and suggests a link between total mitochondrial content of a cell and the nucleoid number.283 nucleoids

56nucleoids

### Nucleoid analysis by mtFociCounter in U2OS cells

To test if *mtFociCounter* is compatible with other cell types, we analysed 220 single U2OS WT cells in three independent experiments (**Figure 4A** and **B**). We detected an average of 221 nucleoids per cell (+/-85 std; n=220 cells) with a coefficient of variation of 38.7%. The average total mitochondrial area of these cells was 194.6μm^2^ (+/-77 std; n=220 cells) with a coefficient of variation of 39.6%, leading to a nucleoid density of 1.2 nucleoids/μm^2^ (+/-0.3 std; n=220 cells) and 22.8% variation, as expected from the preserved correlation between nucleoid numbers and mitochondrial area in (Corr. = 0.79) (**Supplementary Figure 4J**). The nucleoid number and mitochondrial area are both Normally distributed in U2OS WT cells (**Supplementary Figure 4A - I**, p=0.455; Two-sample Kolmogorov-Smirnoff Test).

**Figure 4.**
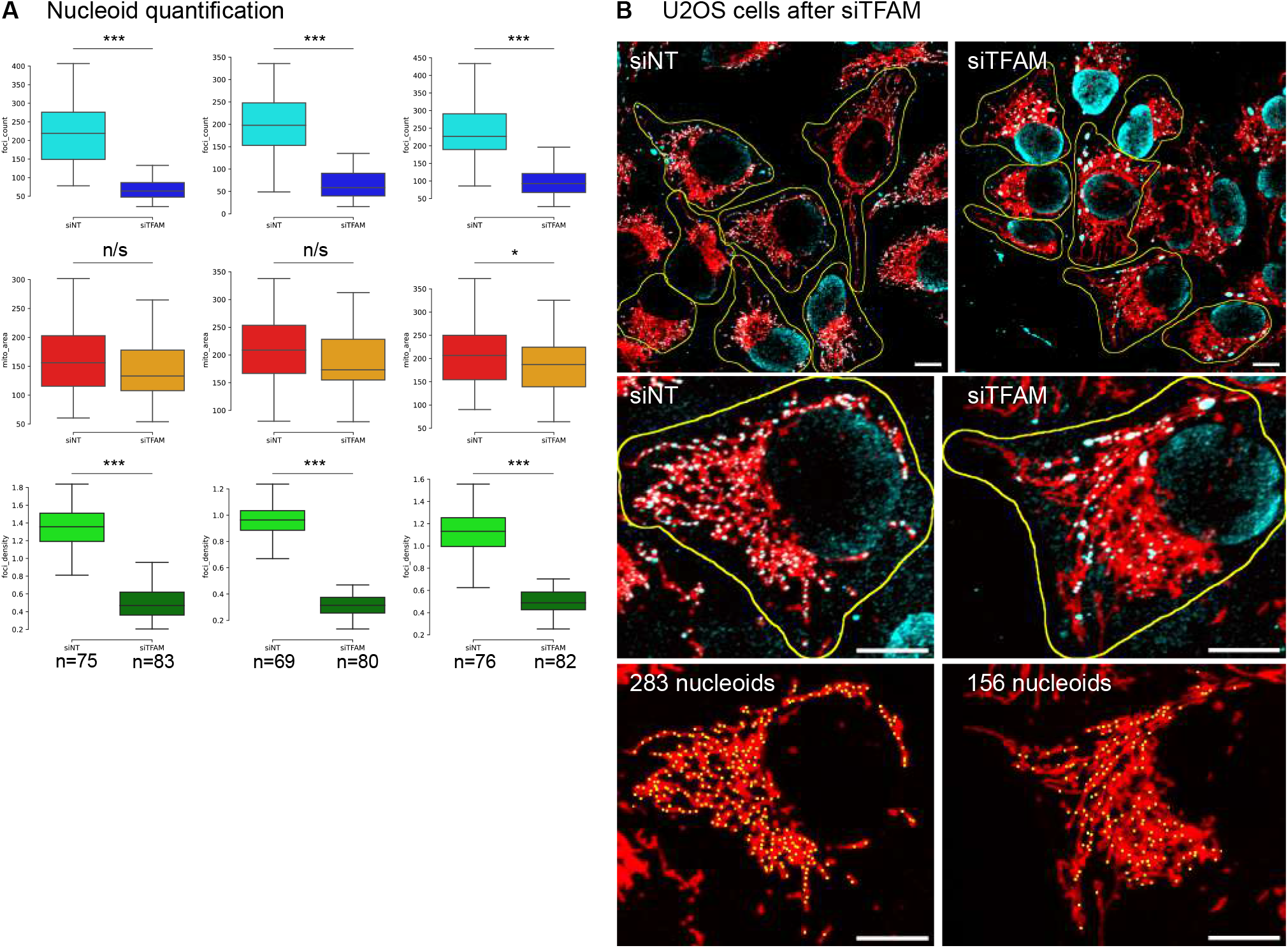
Quantification of nucleoid depletion by *mtFociCounter* in U2OS cells after 3 days of siTFAM treatment. (**A**) Box plots and quantitative analysis for three independent experiments of nucleoid number per cell (siNT = cyan, siTFAM = blue), mitochondrial area in micrometre squared (siNT = red, siTFAM = orange) and nucleoid density per micrometre squared (siNT = lime, siTFAM = green). All box plots represent the median and the first and third quartile. Whiskers represent the rest of the distribution except outliers. Two-sample Kolmogorov-Smirnoff tests were used with alpha = 5% and * denoting p<0.05, *** denoting p<0.0001. (**B**) Representative full FOV of U2OS cells with mitochondria (red) and dsDNA (cyan), treated with neutral siRNA (siNT, left) or siRNA against TFAM (siTFAM, right) and cell segmentations in yellow. Centre with single cell from above, bottom with same cell and detected foci (yellow boxes) using prominence of 82. All scale bars are 10μm.

### mtFociCounter detects siTFAM-induced nucleoid reduction

Next, we intended to verify the performance of *mtFociCounter* in conditions where the mtDNA copy number is perturbed. Knockdown of the mitochondrial nucleoid packaging protein TFAM by siRNA is known to reduce the mtDNA copy-number in HeLa and HEK293 cells [14]. Therefore, we analysed the images of U2OS WT cells treated with siTFAM or non-targeted siRNA controls (siNT) for three days and from three independent experiments by *mtFociCounter* (**Figure 4A** and **B**). We found siTFAM treated U2OS cells have an average of 81 nucleoids per cell (+/-45 std; n=245 cells) with a coefficient of variation of 55.9%. The average total mitochondrial area of siTFAM treated cells was 176.6μm^2^ (+/-65.1 std; n=245 cells) with a coefficient of variation of 36.8%, leading to a nucleoid density of 0.5 nucleoids/μm^2^ (+/-0.2 std; n=245 cells) and 42.1% variation. Interestingly, the nucleoid number and mitochondrial area are borderline to normally distributed in the siTFAM cells (p_nucleoids_=0.00741, p_mitochondria_=0.0506; Two-sample Kolmogorov-Smirnoff Test), and the correlation between nucleoid numbers and mitochondrial area is lower (Corr. = 0.62) (**Supplementary Figure 5A - J**). In direct comparison between the siTFAM and control samples, we thus found non- or barely significant differences between in the mitochondrial area, but robust and stark reduction in the number of nucleoids per cell and nucleoid density in siTFAM treated cells, across all three replicates (**Figure 4A** and **B**). Together, these experiments demonstrate that *mtFociCounter* can be used for multiple cell types as well as to detect differences in nucleoid numbers and nucleoid densities, as shown by the significant reduction of nucleoid numbers in TFAM-silenced U2OS cells. nucleoids 590 nucleoids 943 nucleoids

### Nucleoid analysis by superresolution microscopy

Several mitochondrial foci, including nucleoids, and the mitochondrial network lie at the resolution limit of fluorescence microscopy [2,3,20]. Diffraction limited microscopy, such as confocal, can therefore systematically underestimate the true number of nucleoids per cell. Superresolution microscopy allows to circumvent the diffraction barrier and to investigate mitochondria and their features at greater detail [3]. We therefore performed superresolution microscopy of 3t3 WT fibroblasts by Elyra7 SIM ^2^ and test whether *mtFociCounter* detects an increased number of nucleoids depending on the image modality applied. Indeed, in superresolved images of 3t3 WT fibroblasts we measured a mean number of 644 nucleoids (+/-345 std, n=200 cells) and a median number of 586 nucleoids (inter quartile range of 357) per single cell, with a coefficient of variation of 53.6% (**Figure 5A**). The observed mean mitochondrial area is 109.0μm^2^ (+/-57.4μm^2^ std; n=200 cells) with a coefficient of variation of 52.6% and a median of 98.3 um^2^ (inter quartile range = 60.5μm^2^) (**Figure 5B**). The correlation between mitochondrial area and the number of detected nucleoids in individual cells was well preserved at 0.90, and the mean nucleoid density is 6.1 nucleoids/μm^2^ (+/-1.6 std; n=200 cells), with a median of 5.9 (inter quartile range = 1.8) and coefficient of variation of 26.6% (**Figure 5C, Supplementary Figure 6J**). Both the observed single-cell distribution of the number of nucleoids and the mitochondrial area are borderline significantly different from a Normal distribution (p_nucleoids_=0.0297, p_mitochondria_=0.0221; Two-sample Kolmogorov-Smirnoff Test), whereas the single-cell nucleoid density is normally distributed (p=0.142; Two-sample Kolmogorov-Smirnoff Test)(**Supplementary Figure 6A - I**). Together, these results show that *mtFociCounter* is readily applicable to different imaging modalities and detects a higher number of nucleoids and a smaller mitochondrial network area with increased image resolution, as expected. Notably, the large cell-to-cell heterogeneity of nucleoid numbers and mitochondrial network size in 3t3 WT fibroblasts are well preserved at superresolution, and thus reflect true underlying biological variance. Furthermore, the higher correlation between mitochondrial area and nucleoid numbers together with the reduced variance upon normalisation support the benefit of assessing the nucleoid density, in addition to absolute nucleoid numbers in single cells. Our experiments also show that superresolution microscopy is necessary to obtain highly accurate descriptor values, but that undersampling errors are largely systematic. This allows to robustly detect relative differences between samples by standard confocal microscopy, with sufficiently large sample and effect sizes. The choice of the imaging set-up thus depends on the biological question, and *mtFociCounter* allows to consistently analyse and assess the results from different experimental systems.

**Figure 5.**
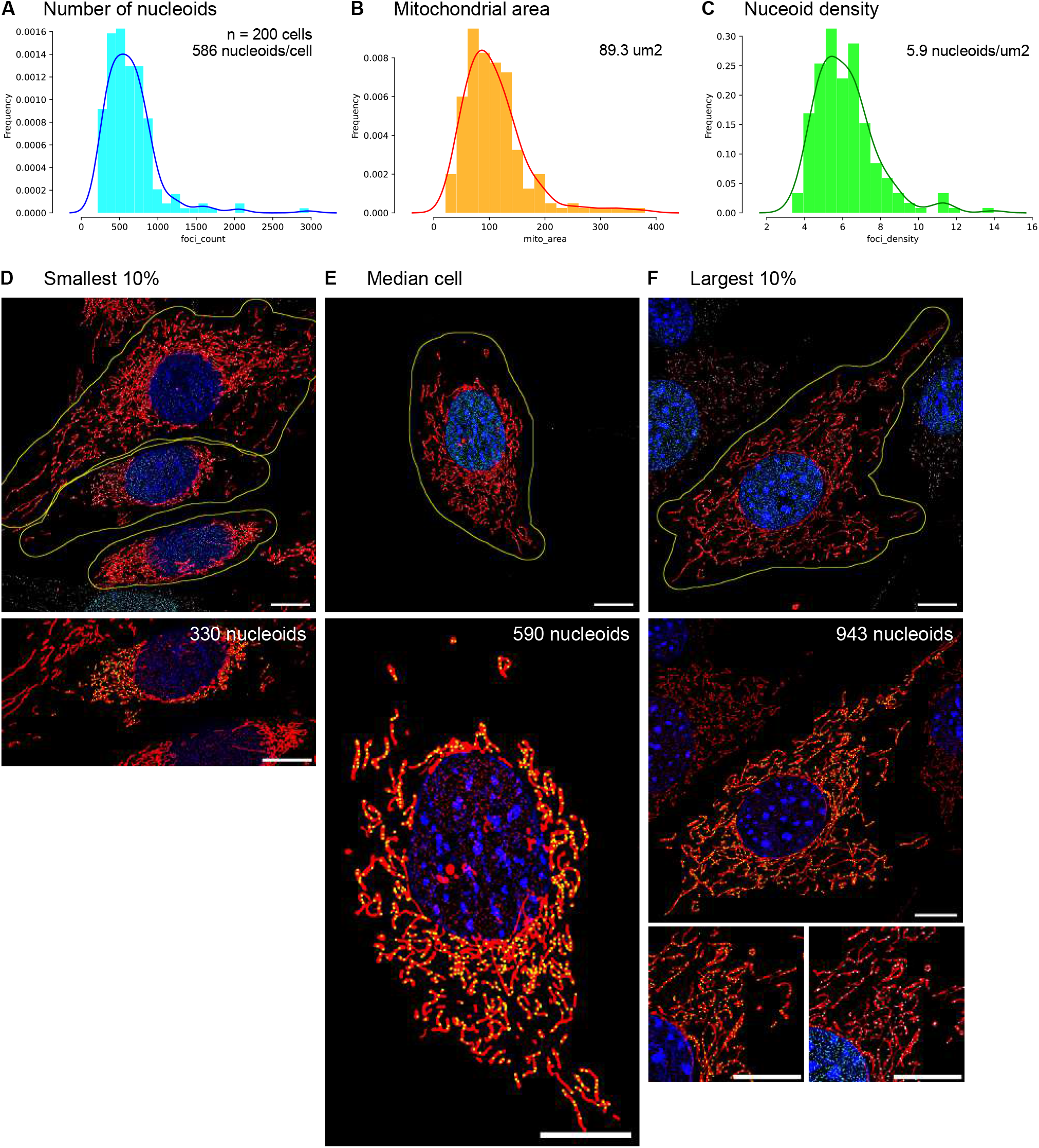
Quantitative analysis of mitochondrial nucleoids in single 3t3 fibroblasts from SIM^2^ superresolution microscopy. (**A**) Histogram and KDE plots of the number of mitochondrial nucleoids, (**B**) the mitochondrial area in square micrometres and (**C**) the nucleoid density (μm^-2^), with median values indicated (n = 200 single cells from 3 independent experiments).(**D**) Full FOV with the cell (centre) denoting the lowest 10% according to the number of detected nucleoids (330), with mitochondria (red), nucleus (blue) and DNA-foci (cyan) and cell-segmentation outline (yellow line) on top or image of the cell with detected foci (yellow boxes) on bottom. (**E**) Full FOV of the cell with the median number of nucleoids (590), and rest as above. (**F**) Full FOV of the cell comprising the 10% highest number of detected nucleoids (943) and as above, with two zoomed excerpts shown below. A prominence level of 800 was used for these analyses. All scale bars are 10μm.

## Discussion

The number of mtDNA molecules and mitochondrial nucleoids in single cells depends on the cell type and cell state [4–7]. Changes in the average copy number per cell have been associated with several diseases [6,8–10], yet the exact mechanisms which determine copy-number homeostasis within cells remain unclear [9]. Today, nucleoids in single cells can be counted manually, by visual inspection [7,22,23], or automatically, with proprietary software [3,24,25]. However, manual counting is typically limited to 10s of cells and both approaches restrict the traceability and reproducibility of analysis and the integration and comparison of results from different experiments. Digital droplet PCR for single-cell mtDNA quantification on the other hand, does not allow to assess contextual information, such as the mitochondrial area per cell [15,16]. Differences in absolute numbers of nucleoids have thus not been differentiated from differences in cell size or cell state occupancy, and previous reports of single cell mtDNA numbers only analysed 10s to few 100s of cells across conditions. In order to investigate the fundamental mechanisms which regulate mitochondrial gene maintenance at the cellular level, simple tools for traceable, reproducible and comparable analysis of mitochondrial nucleoids, capable of preserving context in single cells, are therefore needed today.

*mtFociCounter* is a simple tool for automated quantification of mitochondrial nucleoids, and potentially other foci, from fluorescence images (**Figure 1**). It allows for a reproducible and comparable analysis by generating a perfectly traceable workflow, from raw images to final results. *mtFociCounter* is open source and it can be downloaded and run free of charge [27]. Upon input format conversion, *mtFociCounter* first provides an interface for manual segmentation of single cells from 2D or 3D microscopy images. For foci-detection, the user is then required to perform an automated parameter sweep. We show that this is necessary but sufficient to identify appropriate analysis conditions, avoiding false positives (**Figure 2**). Optionally, detected foci can be filtered by exclusion (“nuclear”) and inclusion (“mitochondrial”) masks, whereby the area of the mitochondrial mask is measured for every cell, in addition to the foci number. The modular design and open source code of *mtFociCounter* allows easy customisation and integration of other software, depending on the experimental set-up and questions, while generating directly comparable output.

To demonstrate the capabilities of *mtFociCounter*, we analyse more than 2’800 single mouse 3t3 fibroblast and human U2OS cells (**Figure 3-5**). We find a very high cell-to-cell variance in the number of nucleoids per cell as well as the mitochondrial area in both cell types, albeit the U2OS cells are slightly less variable. A high variability between cells could be expected from mixed populations of proliferative cells. However, it is interesting to note that there is more than a two-fold increase between the smallest 10% and the largest 10% of cells, indicating the doubling of cellular materials during the cell cycle alone cannot explain the entire variability. In addition, we observe a good correlation between the mitochondrial area and the number of nucleoids in single cells of both cell types (**Supplementary Figure 3** and **4**). Notably, the normalised number of nucleoids per mitochondrial area in single cells, which we term nucleoid density, is less variable than the absolute nucleoid number per cell (**Figure 3-5**). Together, these findings indicate that the absolute number of nucleoids per cell may underly less strict regulation than the relative density of mtDNA per mitochondrial area. *mtFociCounter* will allow to investigate this question on the regulation of cellular mtDNA homeostasis in the future. This example further emphasizes the advantage of context preservation in single-cell analysis, in comparison to bulk-cell assays.

Our superresolution microscopy experiments demonstrate that *mtFociCounter* can be applied to several experimental set-ups, and that the mitochondrial area and exact nucleoid number can be estimated more precisely with increased image resolution. However, superresolution microscopy has several practical disadvantages, throughput limitations and access can be restricted. We find that spinning disk confocal microscopy was sufficient to detected a correlation between nucleoid numbers and mitochondrial area, that the cell-to-cell variability is consistent, and that a clear difference between siTFAM treated cells and control cells is readily detected. Comparison of mitochondrial foci detected by *mtFociCounter* is thus possible from multiple experimental setups, and the ideal approach depends on the biological question.

A main limitation of the presented version of *mtFociCounter* is its reliance on manual cell segmentation. While this bears the advantage of straight-forward input-data selection, it reduces analysis throughput. Furthermore, to minimise the complexity of *mtFociCounter* as a whole, the built-in functionality to create exclusion- (“nucleus”) and inclusion-masks (“mitochondria”) is simplistic. The modular architecture and gateways of *mtFociCounter* however, already allows to use sophisticated algorithms, machine learning and deep learning approaches such as ilastik, weka, cellProfiler or StarDist [29–31,36,37] for fully automated cell and feature segmentation, while preserving the traceability and comparability of the foci analysis. Another challenge in single-cell analysis is the large cell-to-cell variability. We found that experimental replicates can differ significantly for small sample sizes (less than 100 cells)(**Figure 2** and **Supplementary Figure 2**). We therefore propose to emphasise on large number of cells in every condition and that control and treated samples are always processed in parallel to allow direct comparison.

In the future, *mtFociCounter* could be extended to detect more than one foci-type in every cell. This would for instance allow to measure the number of nucleoids and mitochondrial RNA granules within the same cell, or to determine the ratio of replicating nucleoids. Furthermore, additional foci-descriptors, such as foci-size or location, as well as contextual information, such as the cell cycle state, could be implemented, and identification of foci in three dimensions could improve detection accuracy due to overlap.

Taken together, we report the development of *mtFociCounter*, a new and simple approach to quantify mitochondrial nucleoids in single cells. It allows to assess the effects of cellular conditions on the mitochondrial genome and to investigate the underlying mechanisms thereof in a fully traceable and reproducible manner. Other organelles and foci-types can of course also be quantified and we anticipate *mtFociCounter* to serve for a wide community of foci-analysts.

## Data Availability

All data and code are freely accessible on the data-repository Zenodo (DOIs: 10.5281/zenodo.7633157, 10.5281/zenodo.7634535, 10.5281/zenodo.7634645, 10.5281/zenodo.7634604) and Github (https://github.com/TimoHenry) respectively. Please do not hesitate to contact the authors in case of unclarity, questions about the use, suggestions for improvements or any other queries. Please make changes to the source code and submit improvements via Github. Biological material may be available upon reasonable request.

## Funding

This research is funded by EMBO Long Term Fellowship 2021-152 awarded to TR, by UKRI MRC core funding to MM and JP (MC_UU_00028 and MC_UU_00028/5 respectively) and Biotechnology and Biological Sciences Research Council (BBSRC) (BB/W008467/1) to JP and LCT.

## Conflict of interest and disclosure

Michal Minczuk is co-founder and scientific advisor of Pretzel Therapeutics Inc. JP is consulting for Pretzel Therapeutics. TR is employed as a Bioprocess Engineer at LONZA. Beyond this, the authors declare no competing interests.

## Contributions

TR conceived the study, devised the software, performed 3t3 experiments, analysed the data and wrote the manuscript. LCT designed and performed RNAi experiments and contributed to the manuscript. JP supervised LCT and proof-read the manuscript. MM hosted TR and proof-read the manuscript.

## Acknowledgments

We would like to thank Roy Chowdhury for meticulous management of the MBU microscopy facility, Pedro Pinheiro-Silva for WT and Pavel Nash for rho0 NIH/3t3 cell lines, Christoph Gäbelein and Julia Vorholt for stable cell line creation, Benoît Kornmann and Quian Feng for plasmids, and Suliana Manley for support.

## Supplementary Material

**Supplementary Figure 1**. Software-architecture of *mtFociCounter*.

**Supplementary Figure 2**. Experimental set-up and reproducibility analysis. (**A**) Confocal image of 3t3 fibroblasts with immunostaining against dsDNA (cyan) and against TOMM20 (mitochondria, red). (**B**) Confocal image of 3t3 fibroblasts with nuclear staining (Hoecst, blue), cell-outline (WGA-488, magenta) and mitochondria (MTS-dsRed, red). (**C**) Confocal image of 3t3 fibroblasts with immunostaining against dsDNA (cyan), MTS-dsRed (red) and Hoechst (blue) showing an actively dividing cell at the centre left. (**D**) Comparative analysis of 3t3 fibroblasts stained with or without a primary antibody. Left, four images from two representative FOVs from samples stained with (bottom row) or without (top row) primary antibodies. Contrast was manually but equally adjusted to highlight all foci based on “no primary” image (left column) or based on control image (right column). Centre, box and scatter plots of number of foci for each prominence value from a prominence sweep. Total number of cells analysed, from the combination of three independent experiments for each condition, are indicated. Right, box plots for direct comparison between primary-less (no1, blue and orange) and primary-stained (wt, cyan and red) of number of detected foci (left) for two prominence values, 5 or 62, and mitochondrial area. (**E**) As **D** for comparison between 3t3 rho0 cells, lacking mtDNA and 3t3 WT cells with normal mtDNA, both stained with antibodies against dsDNA (cyan). (**F**) Box plots for comparison of the number of nucleoids detected from the same sample after two separate rounds of cell segmentation (n=119 analysed cells after first segmentation and n=128 analysed cells after second segmentation). (**G**) Box plots of number of nucleoids detected in four independent experiments. Number of cells analysed for each experiment are indicated and p-value from Kruskal-Wallis test is indicated. All scale bars are 10μm. All box plots show median, first and third quartiles as well as the remaining distributions as whiskers. Two-sample Kolmogorov-Smirnoff tests were used for **D** to **F**, with alpha = 5% and * denoting p<0.05, ** denoting p<0.001, *** denoting p<0.0001.

**Supplementary Figure 3**. Distribution and correlative analysis of single-cell descriptors for 2165 3t3 WT cells. (**A**) Testing Normality of the data distribution by Q-Q plots for mitochondrial nucleoids, (**B**) mitochondrial area and (**C**) nucleoid density. (**D - F**) Comparison between the observed distribution and a simulated Normal distribution based on the observed mean, variance and sample size by 2-sample Kolmogorov-Smirnoff tests (p values are indicated) and visualisation of the respective Cumulative Density Functions, with simulations in red. Overlay of the observed distribution for the number nucleoids (**G**) (cyan), mitochondrial area (**H**) (orange) and nucleoid density (**I**) (green) and the simulated distributions (red). (**J**) Scatterplot, regression line and histogram with KDE plots for correlation analysis of mitochondrial area and the number of nucleoids detected in single cells. The correlation of 0.81 between mitochondrial nucleoid number and mitochondrial area is indicated. (**K**) Q-Q plot analysis of median number of nucleoids detected in 14 independent experiments. For all plots, the full data set of 2165 cells from 14 independent experiments, as described in **Figure 3**, was analysed.

**Supplementary Figure 4**. Distribution and correlative analysis of single-cell descriptors for 220 U2OS cells treated with neutral siRNA. Analogous to **Supplementary Figure 3**.

**Supplementary Figure 5**. Distribution and correlative analysis of single-cell descriptors for 245 U2OS cells treated with siTFAM. Analogous to **Supplementary Figure 3**.

**Supplementary Figure 6** Distribution and correlative analysis of single-cell descriptors for 200 3t3 WT cells imaged by superresolution microscopy. Analogous to **Supplementary Figure 3**.

For supplementary Figures, please visit: https://doi.org/10.5281/zenodo.7838928

## Notes

https://doi.org/10.5281/zenodo.6962215

https://github.com/TimoHenry/mtFociCounter

https://doi.org/10.5281/zenodo.7838928

### Summary of Updates

The software has been revised and to overcome some bugs. Additional data for 3t3 WT cells and superresolution images were added. Control experiments without primary antibody staining and with rho0 cells were added. Exampifying experiments with U2OS cells treated with siRNA against TFAM or control siRNA were added.

